# The ZMYND8 chromatin factor protects cardiomyocyte identity and function in the mouse heart

**DOI:** 10.1101/2022.10.06.510015

**Authors:** Andrew Kekūpaʻa Knutson, Abigail Avelar, Ralph V. Shohet

**Affiliations:** Center for Cardiovascular Research, John A. Burns School of Medicine, University of Hawaiʻi, Honolulu, HI 96813

## Abstract

Appropriate gene expression within cardiomyocytes is coordinated by chromatin factors and is essential for heart function. We investigated the role of the chromatin reader ZMYND8 in the mouse heart using null and conditional knockouts (*Zmynd8-cKO)*. While full-length *Zmynd8* is not required for cardiomyocyte development, *Zmynd8-cKO* mice develop cardiomegaly, decreased cardiac function, and premature death compared to controls. Transcriptome analysis of *Zmynd8-cKO* cardiomyocytes reveals illegitimate expression of transcripts normally limited to skeletal muscle. Additionally, we observe integration of TNNI2 skeletal troponin into cardiac sarcomeres of mutant mice. We conclude that ZMYND8 is necessary to maintain appropriate cardiomyocyte gene expression and cardiac function.

## INTRODUCTION

Cardiomyocytes are highly specialized, energy-demanding cells that produce contraction of the cardiac chambers. These cells are extremely sensitive to ATP depletion and are often the first to die under conditions of hypoxia (Knutson et al. 2021). Because adult cardiomyocytes cannot replicate, protecting these cells is a crucial goal of modern cardiovascular science. Defining the genetic factors and mechanisms that protect the cardiomyocyte gene program will improve treatments for both congenital and acquired heart disease (Kathiresan and Srivastava 2012).

The role that chromatin regulators play in coordinating cardiomyocyte development and the progression of heart disease is becoming increasingly apparent (Greco and Condorelli 2015). These factors regulate transcription through several mechanisms including deposition of covalent histone marks associated with active or repressive gene expression, recruitment of transcriptional machinery, or enhancer and suppressor activity. A group of chromatin proteins containing bromodomain motifs have recently attracted attention as critical regulators of the progression of illness (Fujisawa and Filippakopoulos 2017).

ZMYND8 (also known as PRKCBP1 or RACK7) was initially characterized as a binding partner of Protein Kinase C in humans (Fossey et al. 2000) and has since been studied for its role in regulating gene expression, enhancer activity, and the DNA damage response (Shen et al. 2016; Spruijt et al. 2016). While ZMYND8 is not predicted to have enzymatic activity, it can recognize covalent modifications on histone tails including H3K4me1 and H3K14ac (and potentially other marks) through its PHD, Bromo, and PWWP domains (Li et al. 2016; Savitsky et al. 2016). ZMYND8 recruits several chromatin writers and erasers to modulate the chromatin landscape. Although the role of ZMYND8 in the heart is unknown, mutations have been identified in genetic studies of atrioventricular septal defects and syndromic congenital cardiac abnormalities also associated with neural and hearing defects (Al Turki et al. 2014; Dias et al. 2022). Our lab has identified a role for ZMYND8 in suppressing the activity of hypoxia inducible factor, HIF-1, in a transgenic mouse model that expresses oxygen-stabilized HIF-1α in cardiomyocytes (Schunke et al. 2019). Interestingly, acetylated ZMYND8 has both activating and repressive roles in HIF-regulated gene expression in human breast cancers (Chen et al. 2018). These observations prompted us to investigate the role of ZMYND8 in mammalian cardiomyocytes.

Here, we describe physiological, cellular, and molecular consequences of removing full-length ZMYND8 from mouse cardiomyocytes. We find that while ZMYND8 is necessary for embryonic development, it is not necessary for cardiomyocyte development using two established Cre recombinase lines. However, adult mice lacking full-length ZMYND8 in their cardiomyocytes develop enlarged hearts, reduced cardiac function, and distorted cardiomyocyte shape compared to control mice. These mice have a shortened lifespan compared to control mice, with a larger effect in female animals. We also show that in young adult hearts, ZMYND8 represses specific genes, some of which are normally restricted to skeletal muscle. These data characterize a mouse model of *Zmynd8-*related cardiomyopathy and reveal roles for ZMYND8 in maintaining cardiac homeostasis.

## RESULTS AND DISCUSSION

### *Zmynd8* is essential for embryogenesis but dispensable for cardiac development

ZMYND8 belongs to a large family of bromodomain-containing proteins that regulate chromatin and gene expression (Fujisawa and Filippakopoulos 2017). These factors have cell-type-specific functions and are important in normal development and the progression of disease. The role of ZMYND8 in the heart is unknown. To test the necessity for ZMYND8 in the heart, we performed crosses using null and conditional alleles of *Zmynd8* (Figure 1). The flox sites for the *Zmynd8*^*fl*^ allele (Delgado-Benito et al. 2018) span exon 4 of *Zmynd8*, which encodes amino acids 34-85 of the full-length protein (Figure 1). These amino acids encode the entire bipartite nuclear localization sequence for ZMYND8. We isolated a recombined allele of *Zmynd8* (i.e., *Zmynd8*^*Δ*^) that could be inherited through the germline of heterozygous mice, thus affecting all cells of the embryo (see Materials and Methods section). No *Zmynd8*^*Δ/Δ*^ mice were born from crosses of *Zmynd8*^*Δ/+*^ heterozygous mice (Figure 1B).

**Figure 1.**
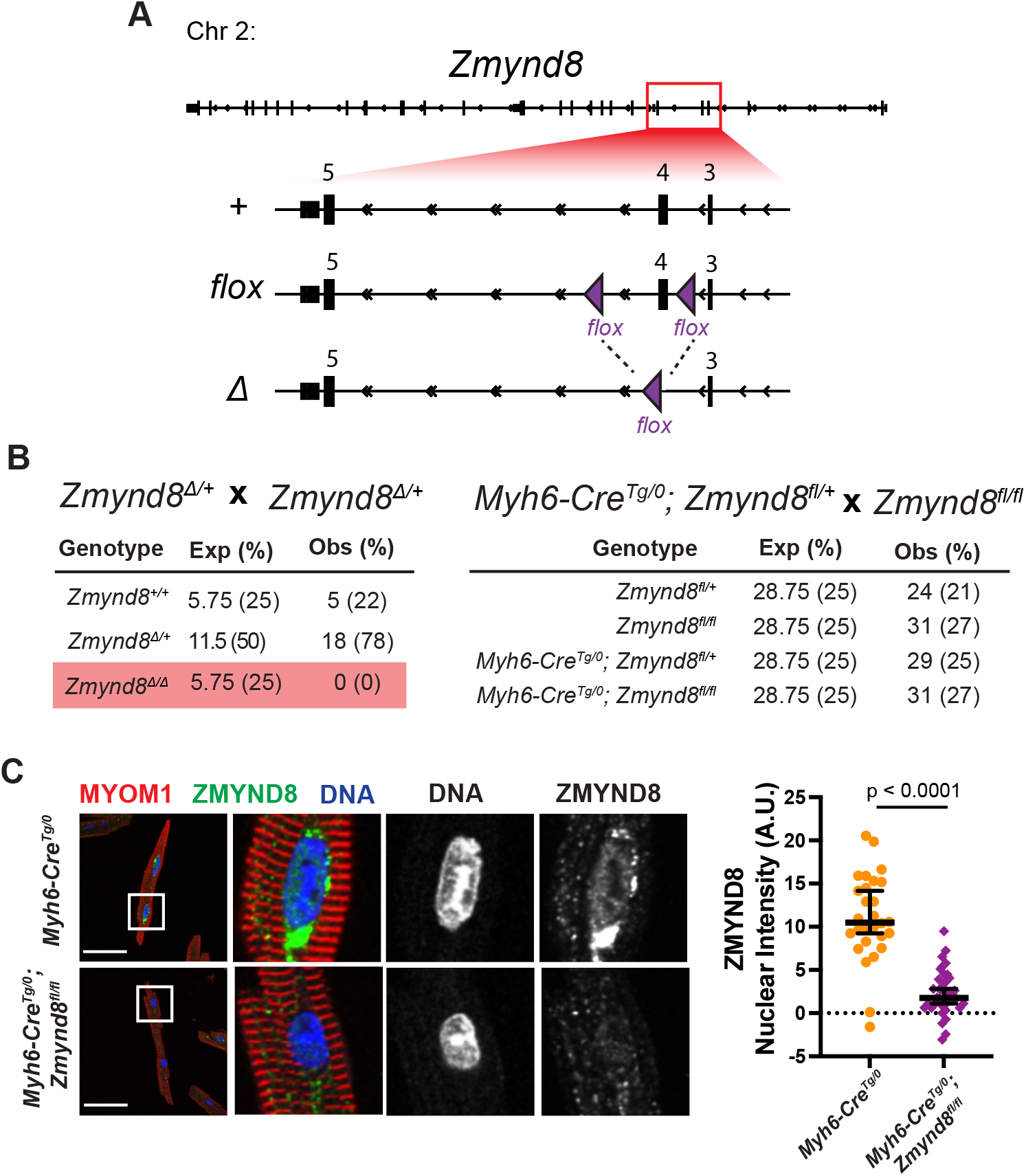
ZMYND8 is necessary for embryonic development but is not required for cardiomyocyte development. **A)** Schematic of the *Zmynd8* gene with the wild-type (+), floxed (flox), and delta (*Δ*) alleles. **B)** Breeding strategy and mendelian ratios of *Zmynd8*^*Δ*^ crosses (left) and *Myh6-Cre*^*Tg/0*^; *Zmynd8*^*fl/fl*^ crosses (right). Expected (Exp) and observed (Obs) number of progeny and percentages are indicated for each cross. **C)** Immunofluorescence staining of control (*Myh6-Cre*^*Tg/0*^) and *Zmynd8-cKO* cardiomyocytes for MYOM1 (red), ZMYND8 (green), and DNA (blue). The nuclei outlined in boxes are expanded to the right. Scale bars are 50 μm. Normalized ZMYND8 nuclear intensity comparisons between control and *Zmynd8-cKO* cardiomyocytes is to the right with statistical significance calculated using an unpaired t-test. Perinuclear accumulation of ZMYND8 is evident in the control cardiomyocyte.

To understand the requirement for ZMYND

8 specifically within the cardiomyocyte lineage, we used two Cre recombinase lines, *Myh6-Cre* (Agah et al. 1997) and *Nkx2*.*5-Cre* (Stanley et al. 2002), in combination with the *Zmynd8*^*fl*^ allele. Progeny from crosses of *Myh6-Cre*^*Tg/0*^; *Zmynd8*^*fl/+*^ males and *Zmynd8*^*fl/fl*^ females were born at expected Mendelian ratios and survived to adulthood (Figure 1B). We repeated these crosses with *Nkx2*.*5-Cre*, which is active in cardiac precursors as early as E7.5 but may also induce recombination in the proepicardium and derivative cell types. Using the *Nkx2*.*5-Cre* line, we also observed progeny born at expected Mendelian ratios (Supplemental Fig. S1A).

We detected efficient flox recombination in heart tissue (Supplemental Fig. S1B) and differences in ZMYND8 protein sizes in *Zmynd8-cKO* hearts compared to control hearts using a monoclonal ZMYND8 antibody (Supplemental Fig. S1C). The observed proteins using this antibody in both control and *Zmynd8-cKO* heart tissue were approximately 30 kD lower than the predicted size of full-length ZMYND8. Compared to control hearts, a larger ∼115 kD band is depleted and a shorter ∼110 kD band becomes enriched in *Zmynd8-cKO* hearts. One explanation for this is that shorter isoforms of ZMYND8 exist in the heart and become preferentially expressed in this model. Consistent with this, immunoblots with a ZMYND8 polyclonal antibody showed several shorter proteins ranging in size from ∼40 to 130 kilodaltons that were detected in both control and *Zmynd8-cKO* hearts (Supplemental Fig. S1C). Full-length ZMYND8 could be detected with this polyclonal antibody from isolated spleens, which contain B cells that are known to express full-length ZMYND8 (Delgado-Benito et al. 2018). Another possibility is that cleavage of ZMYND8 occurs using our isolation methods where removal of exon 4 exposes a proteolytic site that is othewise protected. Indeed, ZMYND8 contains an intrinsically disordered region, which contributes to its ability to aggregate and form liquid condensates at super enhancers (Jia et al. 2021).

To confirm that ZMYND8 was depleted in cardiomyocyte nuclei, we performed immunostaining of isolated cardiomyocytes from *Zmynd8-cKO* mice and observed a marked reduction in ZMYND8 nuclear staining compared to control cardiomyocytes (Figure 1C and Supplemental Fig. S1D).

Interestingly, we sometimes observed strong perinuclear localization in addition to nuclear staining of ZMYND8 in control cardiomyocytes. ZMYND8 has been reported to have both nuclear and cytoplasmic localization in other cell types, including neurons (Yao et al. 2017). These results suggest that while full-length *Zmynd8* is necessary for embryonic development, it is not necessary for cardiomyocyte development, using the Cre lines we tested.

### Knockout of *Zmynd8* in cardiomyocytes leads to cardiac dysfunction and death

While *Myh6-Cre*^*Tg/0*^; *Zmynd8*^*fl/fl*^ mice appeared normal during early adulthood, these mice had a reduced lifespan compared to *Myh6-Cre*^*Tg/0*^ controls with most dying between 5 and 6 months of age (Figure 2A). Signs of distress prior to death included lethargy, tachypnea, a hunched back, and retention of fluid. We observed a difference between male and female mortality with female *Zmynd8-cKO* mice having a slightly shorter lifespan than males (Supplemental Fig. 2). This contrasts with *Myh6-Cre*^*Tg/0*^ control mice where no difference in lifespan between the sexes was detected (Supplemental Fig. 2). Heterozygous mice also showed a sex difference in lifespan with females having a shorter lifespan than males (Supplemental Fig. 2). Additionally, in analyzing male and female *Zmynd8-cKO* siblings born in the same litter, female *Zmynd8-cKO* mice always died before their male siblings. ZMYND8 interacts with the X-linked demethylase KDM5C (Shen et al. 2016) and its Y-linked counterpart KDM5D (Li et al. 2016), raising the possibility that male and female *Zmynd8-*deficient cardiomyocytes manifest sexually-dimorphic responses to stress. KDM5C and KDM5D are thought to be redundant in preventing a compact myocardium phenotype during embryogenesis (Kosugi et al. 2020), however no study has characterized deficiency of these enzymes isolated to the cardiomyocyte lineage or later in life.

**Figure 2.**
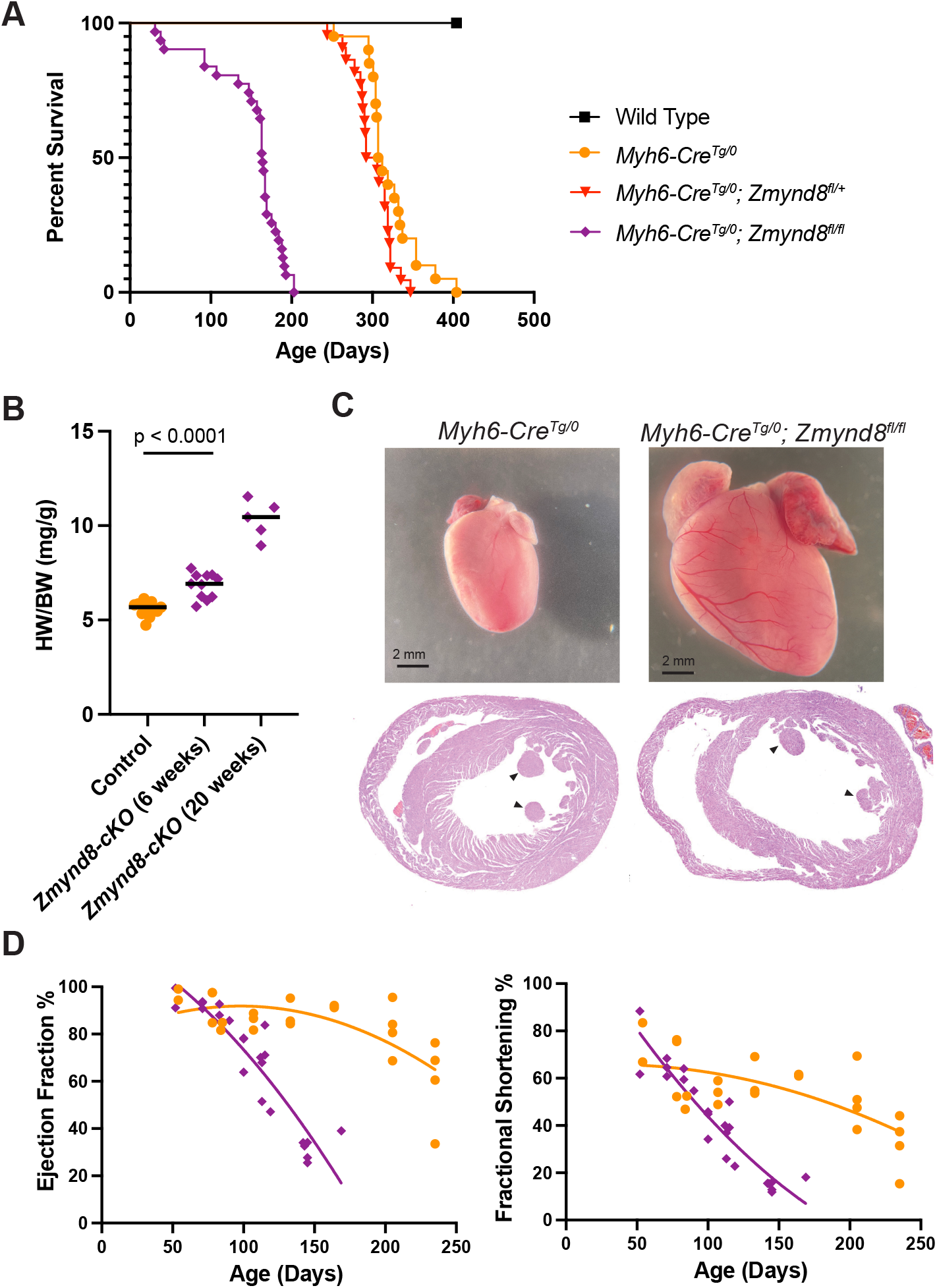
Mice lacking *Zmynd8* in cardiomyocytes die with enlarged hearts. **A)** Lifespan analysis of wild type (black circles), *Myh6-Cre*^*Tg/0*^ (orange circles), *Myh6-Cre*^*Tg/0*^; *Zmynd8*^*fl/+*^ (red triangles); and *Myh6-Cre*^*Tg/0*^; *Zmynd8*^*fl/fl*^ (purple diamonds). Curve comparisons, p-values, and number of animals in each group can be found in Supplementary File 1. **B)** Heart weight to body weight ratios of 6-week old control and 6-week and 20-week old *Myh6-Cre*^*Tg/0*^; *Zmynd8*^*fl/fl*^ mice. **C)** Stereoscopic image of whole hearts from 4-month-old control and *Zmynd8-cKO* male mice (top) and H&E staining of transverse sections (bottom). Arrowheads indicate papillary muscles. **D)** Ejection Fraction and Fractional Shortening measurements of *Myh6-Cre*^*Tg/0*^ control and *Myh6-Cre*^*Tg/0*^; *Zmynd8*^*fl/fl*^ adult mice over time. Each point is an individual mouse with best-fit curves calculated using non-linear regression.

We assessed heart morphology and function in *Zmynd8-cKO* mice and detected an increase in heart weight to body weight ratio as early as 6 weeks compared to sibling controls in both males and females (Figure 2B and 2C, Supplemental Fig. 2D). This heart enlargement progresses over time, with 5-month-old *Myh6-Cre*^*Tg/0*^; *Zmynd8*^*fl/fl*^ mice presenting an increase of 80% in heart mass compared to age-matched control mice (Figure 2B). Cross sections of 8-week-old *Myh6-Cre*^*Tg/0*^; *Zmynd8*^*fl/fl*^ mice showed thinning of the ventricular walls suggesting cardiac remodeling (Figure 2C). We assessed heart function in *Zmynd8-cKO* adult mice using echocardiography and observed a dramatic decrease in contractility over time compared to *Myh6-Cre*^*Tg/0*^ control mice (Figure 2D). These results show that ZMYND8 is necessary to prevent the onset of contractile dysfunction that leads to heart failure and death.

### *Zmynd8-cKO* cardiomyocytes misexpress skeletal muscle transcripts

In addition to its role in regulating enhancer activity (Shen et al. 2016) and DNA damage (Spruijt et al. 2016), ZMYND8 is known to have repressive functions on transcription by recruiting other chromatin factors. To understand the effects of removing *Zmynd8* on cardiomyocyte gene expression, we performed transcriptome analysis of pure populations of rod cardiomyocytes isolated from control and *Zmynd8-cKO* mice (Figure 3A). We assayed young mice (4-weeks old) to capture the primary effects of *Zmynd8* knockout and to avoid secondary transcriptional effects due to heart failure (Figure 2D).

**Figure 3.**
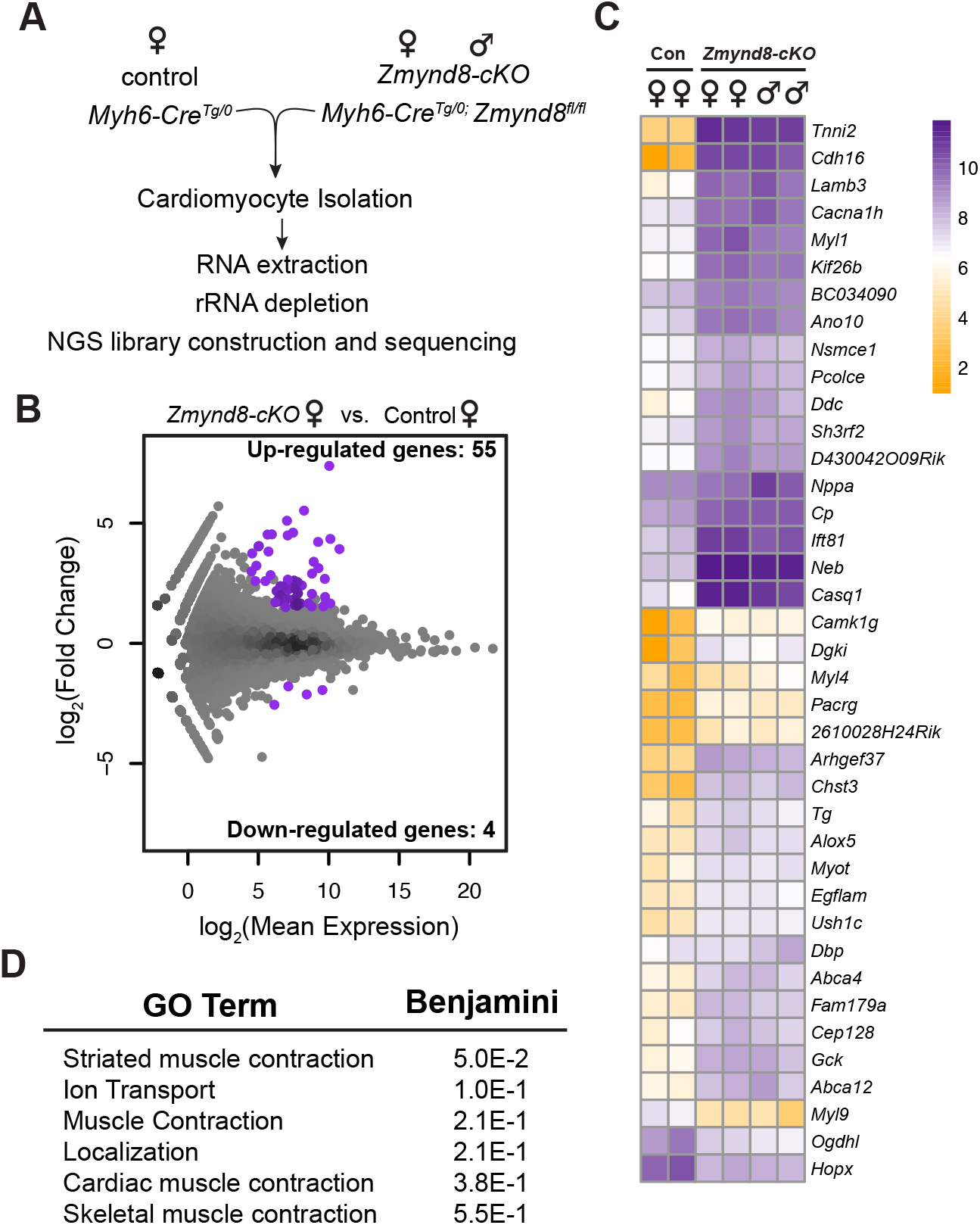
Gene expression changes in *Zmynd8-cKO* adult cardiomyocytes. **A)** Schematic of RNA-seq experiment. Cardiomyocytes were isolated from 4-week-old control (two females) and *Zmynd8-cKO* (two females and two males) mice and RNA was extracted, rRNA depleted, and sequenced. **B)** MA plot of genes up-and down-regulated in 4-week old *Zmynd8-cKO* female cardiomyocytes versus control female cardiomyocytes. Significant genes are colored purple and the number of up- or down-regulated genes are indicated. **C)** Heatmap of normalized log_2_(RPKM) values for each replicate. Genes are listed based on heatmap clustering. **D)** Gene ontology analysis of 55 upregulated genes using DAVID software analysis.

As expected, reads mapping to exon 4 of *Zmynd8* were detected in control samples but were markedly reduced in *Zmynd8-cKO* cardiomyocytes, suggesting both a high degree of cardiomyocyte purity and efficient recombination of the flox sites (Supplemental Fig. 3A). We did find reads mapping to later exons of *Zmynd8* in *Zmynd8-cKO* cardiomyocytes and the level of these reads seemed to increase upon Cre recombination. Regulation of the *Zmynd8* transcript, perhaps through changes in mRNA splicing, or autoregulation of the *Zmynd8* locus by ZMYND8 and its associated machinery, may explain these observations.

We observed subtle changes in gene expression in young cKO cardiomyocytes. Using a cutoff of 1.5-fold in either direction and an adjusted p-value of less than 0.05, we observed 55 genes upregulated and 4 genes downregulated in 4-week post-partum *Zmynd8-cKO* cardiomyocytes (Figure 3B). Gene ontology analysis of the 55 upregulated genes produced categories of striated muscle contraction and genes related to cation transport (Figure 3C and 3D). Compared to control cardiomyocytes, certain genes normally expressed in skeletal muscle are abundant in *Zmynd8-cKO* cardiomyocytes including those encoding the fast-twitch troponin *Tnni2*, the calcium sensor *Casq1*, and the skeletal muscle myosin *Myl1* (Figure 3C and Supplemental Fig. 3). Cardiomyocytes expressing skeletal muscle contractile genes have been observed in embryonic mouse hearts deficient in NuRD complex components including *Hdac1/2* (Montgomery et al. 2007), *Chd4* (Gomez-Del Arco et al. 2016; Wilczewski et al. 2018) as well as adult hearts lacking *Ezh2* (Delgado-Olguin et al. 2012). ZMYND8 interacts with the NuRD subunit GATAD2 (Spruijt et al. 2016), as well as phosphorylated EZH2 (Tang et al. 2021) although these studies were not performed in the heart. Additionally, CHD4 has recently been shown to be recruited by GATA4 and NKX-2.5 to specific loci to repress gene expression programs in embryonic cardiomyocytes (Robbe et al. 2022); it is unknown if ZMYND8 is also involved in this process. In addition to *Chd4, Ezh2*, and now *Zmynd8* mutants, illegitimate expression of skeletal genes in cardiomyocytes has also been observed in mice lacking *Prox1* (Petchey et al. 2014), *Ctcf* (Gomez-Velazquez et al. 2017), the RNA-binding protein TRBP (Ding et al. 2015), or the H3.3 histone chaperone HIRA (Dilg et al. 2016). Defining how these factors work in concert to normally repress skeletal muscle gene expression in cardiomyocytes should be a focus of future work.

Although there were only 4 downregulated genes in *Zmynd8-cKO* cardiomyocytes compared to control they include the cardiomyocyte transcription factor *Hopx* and the cardiac myosin light chain gene *Myl9* (Figure 1C and Supplemental Fig. S3C). HOPX is a homeodomain protein that has been shown to direct cardiomyoblast formation but has not been extensively studied in adult cardiomyocytes (Jain et al. 2015). Based on these data, we conclude that ZMYND8 influences expression of a limited number of important genes in cardiomyocytes. Correct expression or repression of these genes in cardiomyocytes may be necessary for proper cardiomyocyte function.

### ZMYND8 regulates cardiac sarcomere formation and cardiomyocyte morphology

Cardiomyocyte sarcomeres rely on appropriate stoichiometry of their components to efficiently generate contractile force. Abnormalities in sarcomere structure can lead to a range of cardiomyopathies (Lehman et al. 2022) and integration of skeletal troponin can affect cardiac sarcomere contraction (Wilczewski et al. 2018). To determine if skeletal muscle troponin is incorporated into sarcomeres in *Zmynd8-cKO* cardiomyocytes, we performed immunohistochemistry for the cardiac M-band protein MYOM1 and the skeletal troponin TNNI2. We observed robust TNNI2 integration into cardiac sarcomeres in conditional knockout mice using either *Myh6-Cre* or *Nkx2*.*5-Cre* (Figure 3A and Supplemental Fig. S4). Interestingly, only a fraction of cardiomyocytes from each knockout model mis-expressed TNNI2 (Supplemental Fig. S4).

When isolating ventricular cardiomyocytes from *Zmynd8-cKO* mice, we observed that a subset of cardiomyocytes had a distorted shape compared to control. We found that many cardiomyocytes isolated from *Zmynd8-cKO* hearts were shorter and broader than control cells (Figure 4B). We did not observe a correlation between TNNI2 expression and cardiomyocyte shape (Supplemental Fig. S4) suggesting distorted cardiomyocyte morphology is a result of other mechanisms independent of TNNI2 integration into sarcomeres. Taken together, these data show that ZMYND8 regulates the proper formation of cardiac sarcomeres as well as cardiomyocyte morphology in the mouse heart.

**Figure 4.**
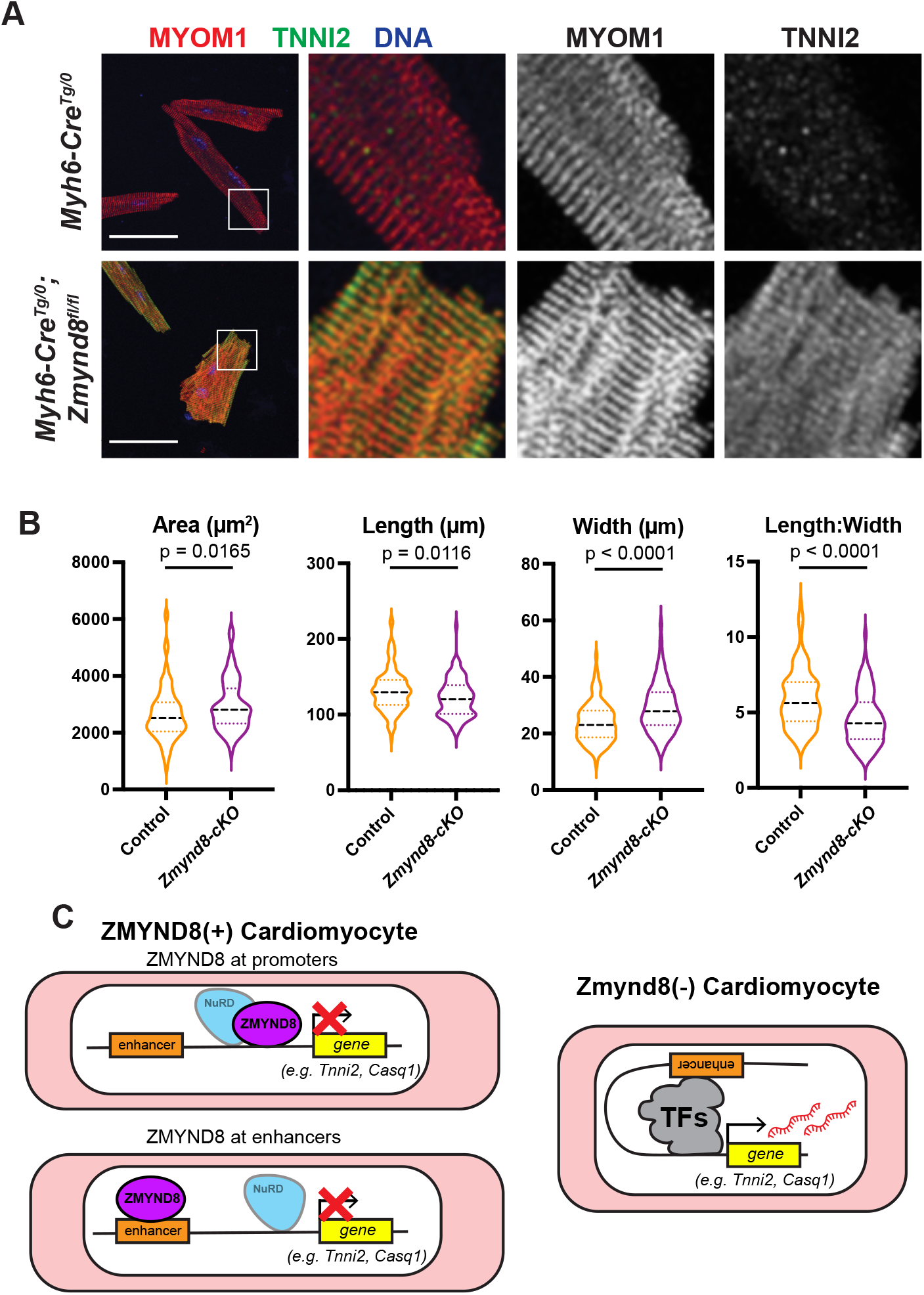
*ZMYND8-cKO* cardiomyocytes integrate TNNI2 into their sarcomeres and have a distorted morphology. **A)** Confocal image of control and *Zmynd8-cKO* cardiomyocytes stained for MYOM1 (red), TNNI2 (green), and DNA (blue). Higher magnifications of boxed region are shown to the right. Scale bars are 50 μm. **C)** Morphometry data for cardiomyocytes isolated from two control (i.e., *Myh6-Cre*^*Tg/0*^) and two *Zmynd8-*cKO adults. At least 75 cardiomyocytes were analyzed from each mouse. P-values are derived from unpaired t-tests. **C)** Models of ZMYND8 activity in cardiomyocytes. ZMYND8 may work with the NuRD complex at promoters to repress gene expression. Alternatively, ZMYND8 may work upstream at enhancer elements, independent of NuRD.

Our study examines the role of ZMYND8 in mouse cardiomyocytes and we believe this work sheds light on the function of this protein in human genetic disease (Al Turki et al. 2014; Dias et al. 2022). Understanding the mechanism by which ZMYND8 prevents cardiac dysfunction will be a focus of future work and could involve ZMYND8 working at promoter and/or enhancers of target genes (Figure 4C). Upon depletion of ZMYND8 from cardiomyocytes, enhancer-promoter interactions at these genes form via other transcription factors, leading to gene expression and the phenotypes we report in this study (Figure 4C). Delineating which factors work with ZMYND8 in cardiomyocytes (e.g., members of the NuRD complex, EZH2, or HIF-1) should reveal valuable insights. Finally, while our experiments correlate ZMYND8’s gene expression and cardiac function, we cannot rule out that the observed physiological phenotypes relate to other functions of ZMYND8 including its role in the DNA damage pathway. Indeed, impaired resolution of DNA damage in cardiomyocytes contributes to progression of cardiomyopathy (Higo et al. 2017). Taken together, our results provide evidence that ZMYND8 is an important chromatin regulator of cardiomyocyte gene expression and cardiac function.

## MATERIALS AND METHODS

### Mouse Strains and Husbandry

*Zmynd8*^*fl*^ mice were generously provided by Dr. Michela Di Virgilo and described previously (Delgado-Benito et al. 2018). This mouse was backcrossed onto a C57BL/6 background for 10 generations before being used in experiments. *Myh6-Cre* JAX#011038 (Agah et al. 1997) and *Nkx2*.*5-*Cre JAX#024637 (Stanley et al. 2002) mice were obtained from Jackson Labs and were backcrossed to C57BL/6 for at least 5 generations before being used in experiments. Mice were maintained in standard conditions with normal chow *ad libitum*. All protocols were approved by the University of Hawaiʻi IACUC. Primers used to genotype the *Zmynd8*^*fl*^ allele and Cre transgenes are in Table S1.

To generate *Zmynd8-cKO* mice, *Myh6-Cre*^*Tg/0*^; *Zmynd8*^*fl/+*^ or *Nkx2*.*5-Cre*^*Tg/0*^; *Zmynd8*^*fl/+*^ males were bred with *Zmynd8*^*fl/fl*^ females. The *Myh6-Cre* transgene, when inherited through the mother, induces recombination in approximately ∼1% of germ cells (Eckardt et al. 2004). This allowed us to isolate the recombined *Zmynd8*^*Δ*^ allele that could be propagated in heterozygotes and confirmed using the *Zmynd8*^*fl*^ primers listed above.

### Echocardiography

Heart function was assessed on unsedated animals using a 32 MHz transducer with a Vevo 2100 system (VisualSonics) as described previously (Williams et al. 2018). Both data collection and analysis were blinded for genotype. Raw echocardiography data are in File S1.

### Cardiomyocyte Isolation

Cardiomyocyte isolation from adult mice were performed using a Langedorff-free method described previously, with minor modifications (Ackers-Johnson et al. 2016). Briefly, after deep anesthesia, the chest was opened and hearts were excised, blood was removed by perfusion and the heart was digested with collagenases 2 and 4 (Worthington) and Protease XIV (Sigma). All buffers were supplemented with 25 nM Blebbistatin (Toronto Chemicals) to inhibit contraction. The atria and great vessels were removed, and ventricular heart tissue was manually dissociated followed by a brief, gentle 2-minute trituration. The resulting cell suspension was filtered through a 100 μm strainer to remove large pieces of tissue. Cardiomyocytes were enriched during gravity settlement steps with gradually introduced Ca^2+^, followed by plating on laminin-coated dishes or coverslips to remove rounded cardiomyocytes and other cell types.

### Immunostaining and Microscopy Analysis

Isolated cardiomyocytes were plated onto coverslips and then fixed in 4% paraformaldehyde for 10 minutes at 37 °C, followed by three washes of PBS at room temperature. Cells were blocked and permeabilized using a PermBlock solution (2% FBS, 2% BSA, and 0.1% IGEPAL CA-630 in PBS) and incubated with primary antibodies overnight at 4 °C. Primary antibodies used for immunostaining include anti-MYOM1 (Developmental Studies Hybridoma Bank), anti-ZMYND8 (Sigma HPA020949), and anti-TNNI2 (Proteintech 11634-1-AP). Cells were washed and incubated with secondary antibodies conjugated with either Alexa Fluor 488 or 594 (ThermoFisher), washed again in PBS-Tween and sealed with Prolong mountant with DAPI. Images were obtained on a Leica SP5 Confocal Microscope with control and experimental images taken at the same laser intensity and exposure. Images were processed using ImageJ and control and experimental images treated identically. To assess nuclear intensity of ZMYND8, we used the polyclonal anti-ZMYND8 antibody.

Briefly, single plane slices of nuclei were imaged and regions of interests (ROI) were determined manually in ImageJ using the DAPI channel. ROI’s were then overlayed onto the ZMYND8 channel and pixel intensities were recorded. To account for background staining, normalization ROIs were defined just adjacent to each nucleus (i.e, in the cytoplasmic region) and subtracted from its corresponding nucleus. The corrected mean nuclear intensities were then plotted.

### Immunoblotting

Immunoblotting of heart tissue has been described previously (Williams et al. 2018). Briefly, heart tissue was pulverized in liquid nitrogen and extracted in RIPA Buffer with protease inhibitors (Roche). Blots were incubated with anti-ZMYND8 monocolonal (Abcam ab201452) or anti-ZMYND8 polyclonal (Sigma HPA020949) antibodies overnight. The next day, blots were washed in PBS + 0.2% Tween-20 and then incubated with fluorescent-conjugated IRDye secondary antibodies (LiCOR). Blots were imaged on a LiCOR Odyssey IR Imaging System.

### RNA-seq and Analysis

Total RNA from isolated cardiomyocytes was extracted in Trizol and only RNA with an RNA Integrity Number of >7 on a Bioanalyzer was used in library construction. Approximately 250 ng of RNA was used to construct each library. Ribosomal RNA was depleted using NEBNext rRNA Depletion Kit original or v2 (Human/Mouse/Rat) (E6310 or E7400) and libraries were built using the NEBNext Ultra II RNA Library Prep Kit for Illumina (E7770). The libraries were multiplexed and sequenced on a NextSeq500 at 75 bp single-end reads.

Reads were aligned to the mm10 mouse genome using HISAT2 (Kim et al. 2019) and duplicate reads were excluded from analyses. Differential gene expression was determined using DESeq2 (Love et al. 2014) with default parameters. Subsequent analyses and plot generation was performed in R. Normalized read counts per transcript (RPKM) values were calculated using CuffDiff2. Reads were converted to bam files and visualized using IGV. Gene ontology analysis was performed using DAVID software. RNA-seq data have been deposited in Gene Expression Omnibus under accession number GSE216254. Processed RNA-seq data are in File S2.

## ACKNOWLEDGEMENTS

We thank Dr. Michela Di Virgilio for kindly sharing the *Zmynd8*^*fl*^ mouse. We also thank Dr. Kathryn Schunke and Dr. Chad Walton for their insights prior to this work being initiated. This work used the Genomics Shared Resource Core at the UH Cancer Center (P30CA071789) the JABSOM Histology Core, and the Mouse Phenotyping Core. Dr. Knutson has been supported by a Ruth R. Kirschstein Postdoctoral Fellowship (F32HL149319), a MOSAIC Career Development Award (K99GM145410), a Burroughs Wellcome Fund Postdoctoral Enrichment Program Award, and a T32 training grant (T32HL115505). A.K.K. and R.V.S designed the experiments.

A.K.K. performed all experiments and analyses except for echocardiography experiments, which were performed and evaluated by A.A. A.K.K. and R.V.S. interpreted data and wrote the manuscript. All authors edited and approved the final manuscript.

