## Supplemental Tables and Figures for "The ZMYND8 chromatin factor protects cardiomyocyte identity and function in the mouse heart"

### TABLE S1. Genotyping primers used in this study

| | Primer Name | Primer Sequence |
|---|---|---|
| *Zmynd8*$^{fl}$ | Prkcbp1_35576_5 | 5’-GACCACAGCTCTTGCACAGG-3’ |
| | ZMYND8_R2(Rev) | 5’-AAGAAAACCCTGAGACCACC-3’ |
| | MDV_p240 | 5’-GTGCAAACGTGTTCAGTGG-3’ |
| *Myh6-Cre*$^{Tg/0}$ and *Nkx2.5-Cre*$^{Tg/0}$ | Cre_F | 5’-TCCAATTTACTGACCGTACACCAA-3’ |
| | Cre_R | 5’-CCTGATCCTGGCAATTTCGGCTA-3’ |

**A**

*Nkx2.5-Cre$^{Tg/0}$; Zmynd8$^{fl/+}$* **X** *Zmynd8$^{fl/fl}$*

| Genotype | Expected (%) | Observed (%) |
|---|---|---|
| *Zmynd8$^{+/fl}$* | 5.75 (25) | 7 (30) |
| *Zmynd8$^{fl/fl}$* | 5.75 (25) | 3 (13) |
| *Nkx2.5-Cre$^{Tg/0}$; Zmynd8$^{fl/+}$* | 5.75 (25) | 7 (30) |
| *Nkx2.5-Cre$^{Tg/0}$; Zmynd8$^{fl/fl}$* | 5.75 (25) | 6 (26) |

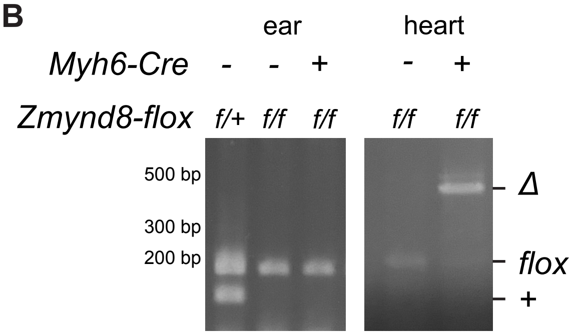


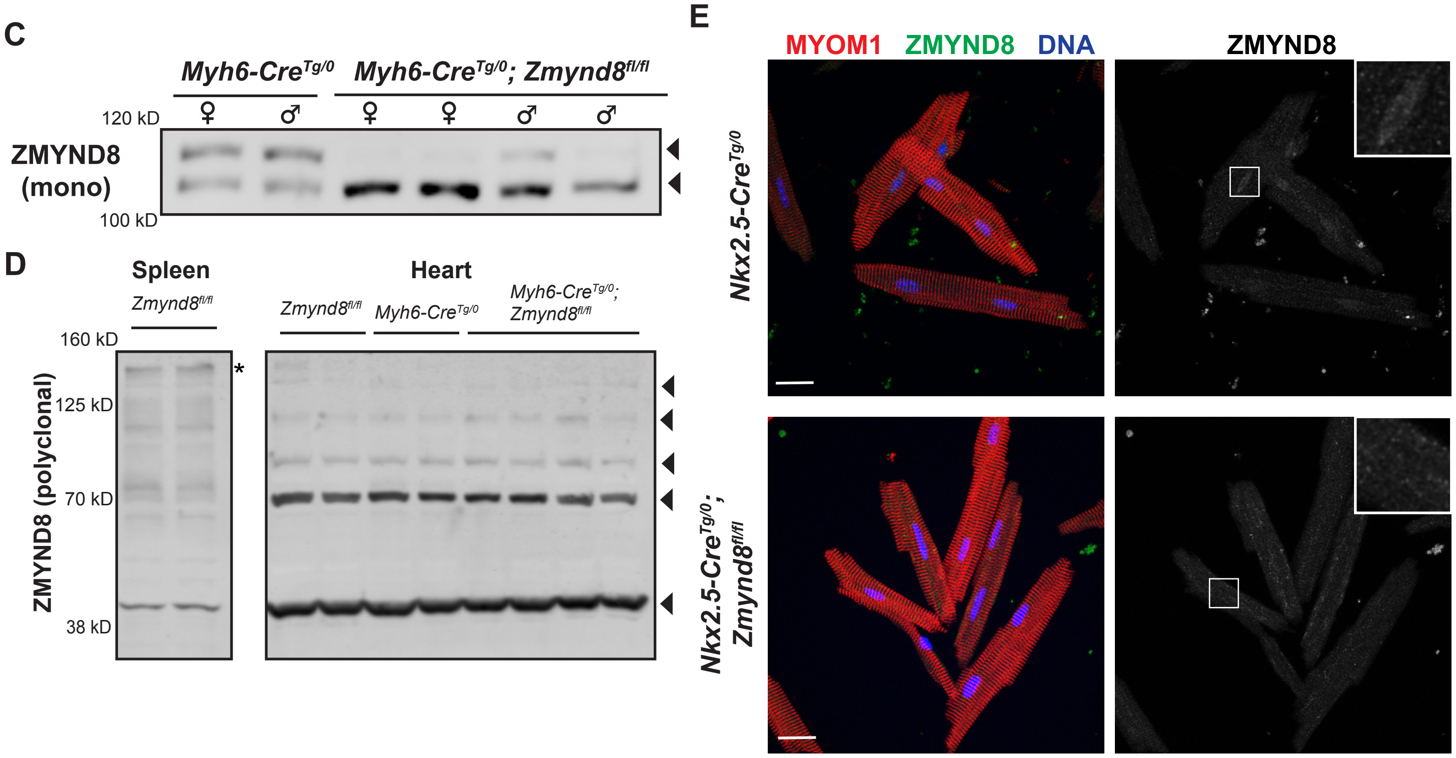


**Supplemental Figure 1.** Conditional knockout of *Zmynd8* from mouse cardiomyocytes. A) Mendelian ratios of F1 progeny from *Nkx2.5-Cre$^{Tg/0}$; Zmynd8$^{fl/+}$* males mated to *Zmynd8$^{fl/fl}$* females. B) PCR amplifcation of ear and heart tissue from *Zmynd8$^{fl/+}$*, *Zmynd8$^{fl/fl}$*, and *Myh6-Cre$^{Tg/0}$; Zmynd8$^{fl/fl}$* mice. The delta (*Δ*), flox, and wild type (+) alleles are indicated to the right. C) Immunoblot of heart extracts from *Myh6-Cre$^{Tg/0}$* and *Myh6-Cre$^{Tg/0}$; Zmynd8$^{fl/fl}$* male and female mice using a monoclonal ZMYND8 antibody. D) Immunoblot of spleen and heart extracts from *Myh6-Cre$^{Tg/0}$* and *Myh6-Cre$^{Tg/0}$; Zmynd8$^{fl/fl}$* mice using a polyclonal ZMYND8 antibody. Asterisk indicates expected size of full-length ZMYND8. E) Immunofluoresence images of cardiomyocytes isolated from *Nkx2.5-Cre$^{Tg/0}$* and *Nkx2.5-Cre$^{Tg/0}$; Zmynd8$^{fl/fl}$* mice stained for MYOM1 (red), ZMYND8 (green), and DNA (blue). Boxed nuclei are expanded in the upper right corner. Scale bars are 20 μm.

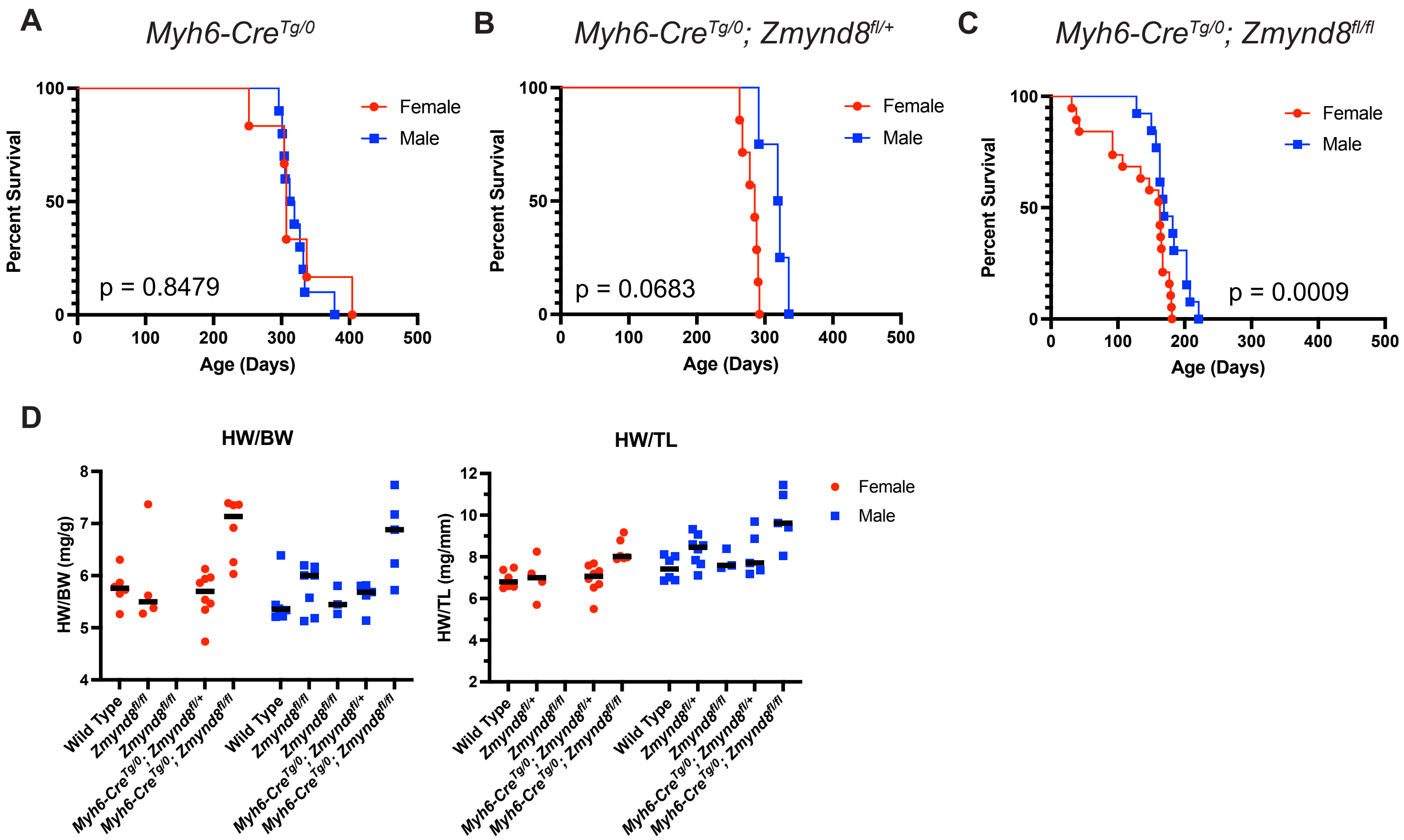


**Supplemental Figure 2.** Lifespan and heart weight measurements disaggregated by sex **A-C)** Lifespan curves for *Myh6-Cre*$^{Tg/0}$ (A), *Myh6-Cre*$^{Tg/0}$*; Zmynd8*$^{fl/+}$ (B) and *Myh6-Cre*$^{Tg/0}$*; Zmynd8*$^{fl/fl}$ (C) mice separated by females (red circles) and males (blue squares). P-values determined by the log-rank test are indicated for each curve. Lifespan comparison data are found in Supplementary File 1. D) Heart weight to body weight or tibia length measurements of male and female 6-week-old mice.

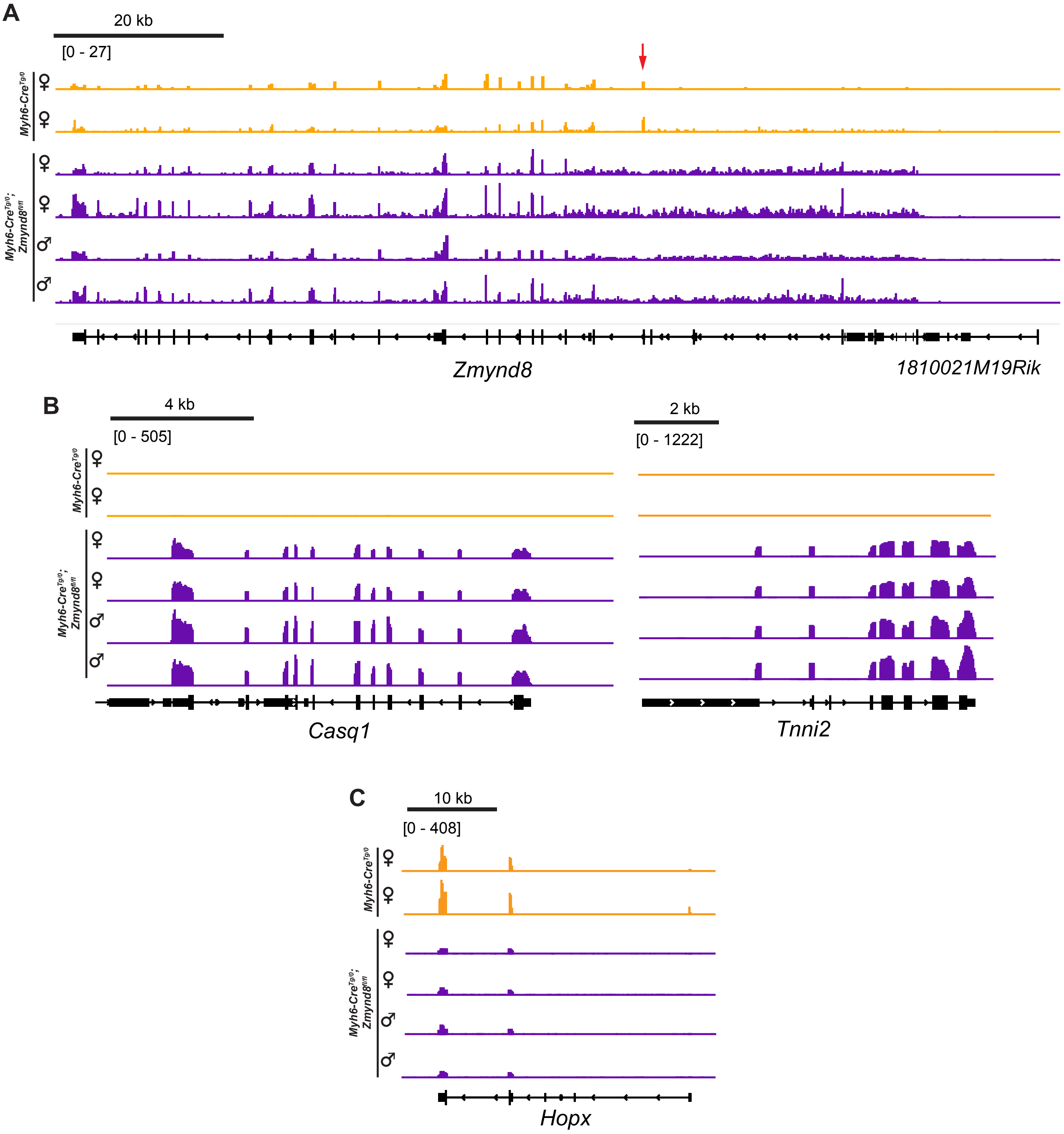


**Supplemental Figure 3.** IGV genome browser tracks showing reads from *Myh6-Cre*$^{Tg/0}$ control (orange) and *Myh6-Cre*$^{Tg/0}$*; Zmynd8*$^{fl/fl}$ (purple) cardiomyocytes across A) *Zmynd8* with exon 4 indicated (red arrow), B) the skeletal muscle genes *Tnni2* and *Casq1*, and C) the homeodomain factor gene *Hopx*. For each gene, all samples are set to the same scale, indicated in brackets below the scale bar.

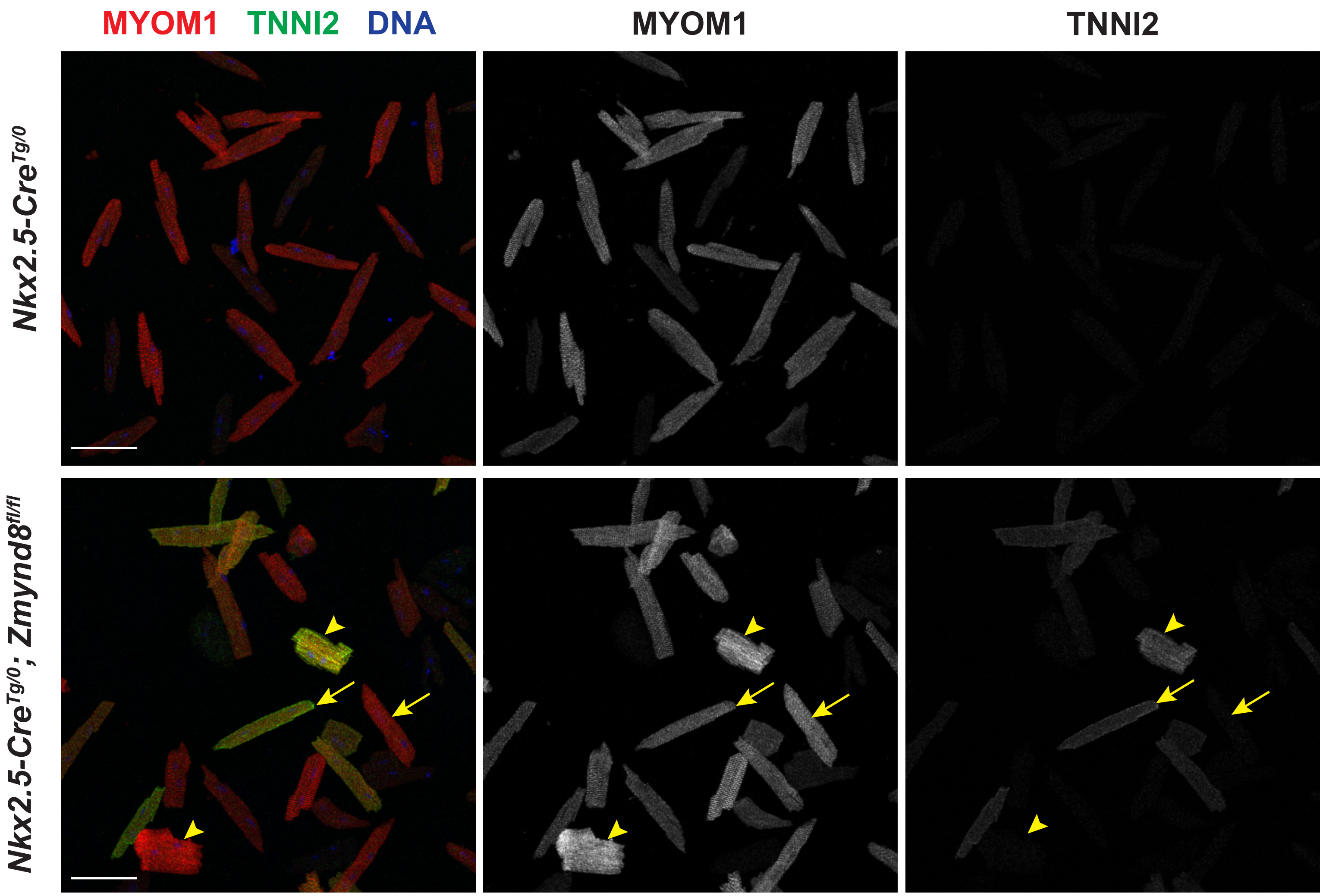


**Supplemental Figure 4.** Cardiomyocytes from *Nkx2.5-Cre*$^{Tg/0}$ or *Nkx2.5-Cre*$^{Tg/0}$*; Zmynd8*$^{fl/fl}$ mice stained for MYOM1 (red), TNNI2 (green), and DAPI (blue). Arrows indicate cardiomyocytes with normal morphology and arrowheads are cardiomyocytes with distorted morphology. Scale bars are 50 μm.
